# Phospholipase Cγ regulates lacrimal gland branching by competing with PI3K in phosphoinositide metabolism

**DOI:** 10.1101/2024.06.28.601066

**Authors:** Qian Wang, Chenqi Tao, Yihua Wu, Karen E. Anderson, Abdul Hannan, Chyuan-sheng Lin, Phillip Hawkins, Len Stephens, Xin Zhang

## Abstract

Although the regulation of branching morphogenesis by spatially distributed cues is well established, the role of intracellular signaling in determining the branching pattern remains poorly understood. In this study, we investigated the regulation and function of phospholipase C gamma (PLCγ) in Fibroblast Growth Factor (FGF) signaling in lacrimal gland development. We showed that deletion of *PLCγ1* in the lacrimal gland epithelium leads to ectopic branching and acinar hyperplasia, which was phenocopied by either mutating the PLCγ1 binding site on Fgfr2 or disabling any of its SH2 domains. *PLCγ1* inactivation did not change the level of Fgfr2 or affect MAPK signaling, but instead led to sustained AKT phosphorylation due to increased PIP3 production. Consistent with this, *PLCγ1* mutant phenotype can be reproduced by elevation of PI3K signaling in *Pten* knockout and attenuated by blocking AKT signaling. This study demonstrated that PLCγ modulates PI3K signaling by shifting phosphoinositide metabolism, revealing an important role of signaling dynamics in conjunction with spatial cues in shaping branching morphogenesis.

## Introduction

Fibroblast growth factor (FGF) signaling is a highly adaptive pathway in biology, influencing a vast array of biological processes from embryonic development and tissue repair to the pathogenesis of various diseases. This remarkable versatility stems from its intricate network of ligands, receptors, co-receptors, and a complex web of downstream signaling pathways complex (Beenken and Mohammadi, 2009; Brewer et al., 2016; Eswarakumar et al., 2005). These components orchestrate a symphony of activation, feedback loops, and crosstalk with other signaling cascades, allowing FGF signaling to exert its diverse effects with exquisite precision. The core FGF signaling pathways include MAPK, PI3K, and PLCγ signaling cascades. MAPK activation originates from the binding of the juxtamembrane region of FGF receptors by the adaptor protein Frs2, which recruits Grb2 and Shp2, triggering Ras-MAPK signaling (Hadari et al., 2001; Ong et al., 2000). Our genetic studies in mice have demonstrated that the adaptor proteins Crk and CrkL, once thought to bind directly to FGF receptors, are also recruited via Frs2 to enhance Ras signaling (Collins et al., 2018). Additionally, while earlier research suggested that PI3K is activated by Gab1 bound to Grb2, our findings reveal that Gab1 is dispensable for FGF signaling (Li et al., 2014). Instead, PI3K signaling is activated by the direct binding of the p110 subunit of PI3K to Ras (Wang et al., 2021). These findings underscore the complex interaction within the FGF signaling network at the cellular level, highlighting the intricate mechanisms that govern its myriad functions.

Compared to Ras and PI3K, the mechanism and role of PLCγ pathway in FGF signaling is much less understood. PLCγ, a member of the mammalian Phospholipase C (PLC) enzyme family, converts phosphatidylinositol 4,5-bisphosphate (PIP2) into diacylglycerol (DAG) and inositol 1,4,5-trisphosphate (IP3) (Kadamur and Ross, 2013). These lipid second messengers trigger the activation of PKC and intercellular calcium release, leading to a plethora of downstream signaling and cellular changes. Unlike most PLC enzymes which are activated by G protein-coupled receptors, PLCγ1 and its relative PLCγ2 are uniquely activated by Receptor Tyrosine Kinases (RTKs) (Yang et al., 2012). Initial biochemical and structural studies demonstrated the N-terminal SH2 domain (nSH2) of PLCγ1 binds the conserved phosphotyrosine residue in FGFR (pY766 in FGFR1) (Bae et al., 2009; Mohammadi et al., 1991), while the C-terminal SH2 (cSH2) domain interacts with the catalytic core of PLCγ1, inhibiting its activity until FGFR phosphorylates tyrosine 783 residue of PLCγ1 to attract binding of cSH2 (Bunney et al., 2012; Hajicek et al., 2019). However, this model was contested by later crystallography studies showing that FGFR2 is bound by the cSH2 domain of PLCγ1, while the nSH2 is dispensable for PLCγ1 enzymatic activity (Huang et al., 2016). The role of PLCγ in FGF signaling is also debated. PLCγ was thought to play a positive role in Ras-MAPK signaling by activating PKC, which can phosphorylate Raf and other MAPK regulatory proteins to stimulate ERK signaling signaling (Lorenz et al., 2003; Marais et al., 1998). Conversely, cell culture studies have shown that FGFR1 lacking the PLCγ-binding residue Y766 displayed reduced internalization (Sorokin et al., 1994), suggesting that PLCγ may promote endocytosis of FGF receptors as a negative feedback mechanism to decrease surface receptor levels. Thus, whether PLCγ exerts a positive or negative effect on FGF-MAPK signaling remains uncertain.

Mammalian lacrimal gland development illustrates the precise regulation by FGF signaling (Garg and Zhang, 2017). The process initiates when the periocular mesenchyme releases Fgf10, which binds to the receptor Fgfr2 on the surface ectoderm, leading to the formation of a lacrimal gland bud followed by an elongated stalk. The bud grows caudally past the junction of supraorbital and infraorbital branches of the stapedial artery until it reaches the final position adjacent to the pinna, where it branches extensively into a tree-like structure (Fig. 1A) (Dartt, 2009; Zoukhri, 2010). Previous studies have suggested that the decision between branching and elongation is influenced by the spatial distribution of FGF, governed by its interaction with heparan sulfate proteoglycans (HSPGs) in the extracellular matrix (Makarenkova et al., 2009; Qu et al., 2012). Strong FGF-HSPG binding leads to a sharp gradient that promotes elongation; weaker binding encourages branching, and a lack of binding results in uncontrolled FGF diffusion, which fails to initiate budding. Interestingly, mutations affecting Fgf10’s affinity to its receptor impact the extent of growth, but not the type of response (Makarenkova *et al*., 2009). This is further supported by observations that reduced lacrimal gland growth results from either a heterozygous mutation in *Fgf10*, increased activity of the negative Ras regulator Sprouty due to Shp2 absence, or deletion of MAPK downstream ETV transcription factors (Garg et al., 2018; Pan et al., 2010; Qu et al., 2011) On the other hand, we recently showed that PI3K signaling is required to prevent the expansion EGF signaling from the lacrimal gland stalk to the bud region, indicating its role in spatial patterning of lacrimal glands (Wang *et al*., 2021). However, the question remains whether intracellular FGF signaling might also determine the choice between branching and elongation in lacrimal gland development.

**Figure 1.**
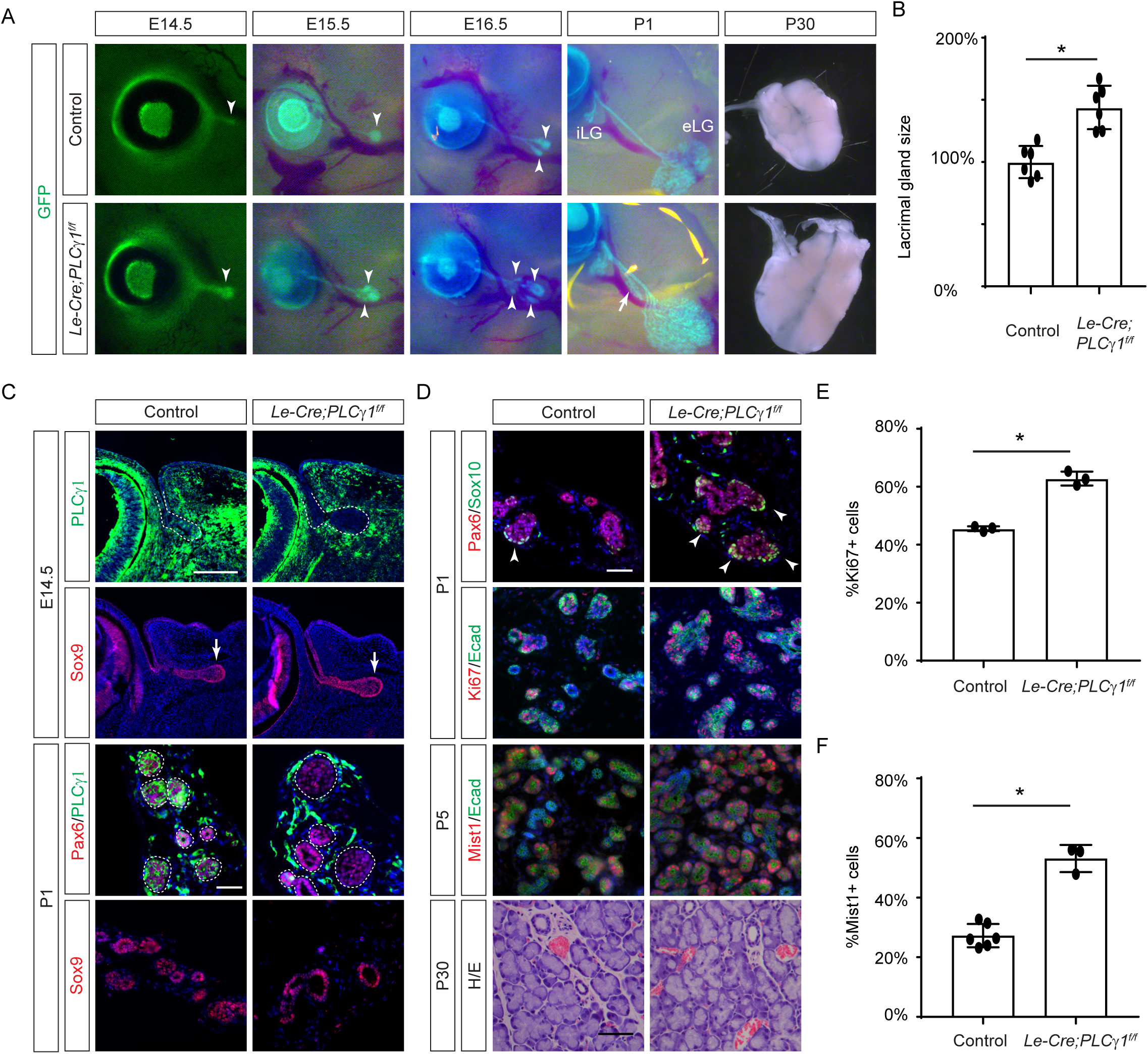
Lacrimal gland hyperplasia in *PLCγ1* conditional knockout mouse. **(A)** Co-expression of the GFP reporter with the *Le-Cre* transgene illustrates the extension of the lacrimal gland bud during embryonic development and its subsequent branching into a glandular structure post-birth. The deletion of *PLCγ1* resulted in extra buds from E15.5 to E16.5 (arrowheads), ectopic branches at P1 (arrow), and enlarged lacrimal glands at P30. **(B)** Quantification of lacrimal gland size at P30. Student’s t-test, *n*=6, *P<0.0001. **(C)** Immunostaining revealed the absence of PLCγ1 in the lacrimal gland bud at E14.5 and the lacrimal gland epithelium at P1 (dotted lines), while the expression levels of progenitor cell markers Pax6 and Sox9 remained unaffected. **(D)** At P1, *PLCγ1*-deficient glands exhibited a rise in Sox10 and Ki67 positive cells, contributing to an increased number of Mist1-expressing acinar cells. By P30, these glands were densely packed with diminished interstitial space. **(E)** Quantification of Ki67-expressing cells. Student’s t-test, *n*=3, *P<0.001. **(F)** Quantification of Mist1-expressing cells. Student’s t-test, *n*>3, *P<0.0001. Scale bars: 50 µm.

In this study, we discovered that *PLCγ* deficiency resulted in hyperplastic lacrimal glands characterized by ectopic branching. This unique phenotype was phenocopied by mutating either the PLCγ binding site on Fgfr2, or any of the SH2 domains of PLCγ1, demonstrating that both SH2 domains are critical for PLCγ activation by FGF signaling. Notably, we observed no significant alterations in Fgfr2 levels, nor was the *PLCγ1* mutant phenotype attenuated upon *Fgfr2* dosage reduction. Furthermore, expression analyses and genetic interaction studies did not implicate the involvement of the MAPK signaling in the *PLCγ1* mutant phenotype. Instead, biochemical and mass spectrometry analyses revealed that FGF-induced AKT phosphorylation persisted in *PLCγ* mutant cells due to a prolonged accumulation in phosphatidylinositol 3,4,5-trisphosphate (PIP3). *Pten* deletion, which promotes AKT activity, exacerbated ectopic branching when combined with *PLCγ1* mutation. Conversely, AKT inhibition curtailed lacrimal gland branching in explants and mitigated the *PLCγ1* phenotype in vivo. These findings collectively demonstrate that PLCγ regulates branching morphogenesis by modulating phosphoinositide metabolism to sharpen FGF-induced AKT signaling.

## Results

### Genetic deletion of PLCγ1 leads to lacrimal gland hyperplasia

To investigate the role of PLCγ in FGF signaling, we generated a conditional knockout of *PLCγ1* using *Le-cre*, which is active in lacrimal gland progenitors (Pan et al., 2008). GFP expression from the *Le-Cre* transgene indicated that *Le-Cre*;*PLCγ1^flox/flox^* mutant lacrimal gland buds were enlarged at E14.5 and exhibited increased branching from E15.5 to E16.5 (Fig. 1A, arrowheads). By postnatal day 1 (P1), while the control animals developed a typical intraorbital gland (iLG) and an extraorbital gland (eLG) linked by a single duct, the *PLCγ1* mutants exhibited not only larger glands but also abnormal branching within the primary duct (Fig. 1A, arrow). By P30, mutant glands were significantly larger than controls, indicating that PLCγ1 likely acts as a negative regulator of lacrimal gland branching morphogenesis (Fig. 1B).

Further molecular analysis revealed that the absence of PLCγ1 immunostaining in *PLCγ1* mutants coincided with enlarged gland buds at E14.5 and expanded clusters of Pax6-positive epithelial cells by P1, aligning with an increased number of Ki67-positive cells (Fig. 1C and E, dotted lines). Although the level of multipotent progenitor markers Pax6 and Sox9 remained unchanged at P1, there were more Sox10-expressing cells in the lacrimal gland end bud, which are known to give rise to acinar and myoepithelial cells (Fig. 1C and D, arrows and arrowheads) (Farmer et al., 2017). This led to an increased number of Mist1-expressing acinar cells by P5 and overcrowded lacrimal gland with little interstitial space by P30 (Fig. 1D and F). These results demonstrate that deletion of PLCγ1 resulted in overgrowth of the lacrimal gland.

### PLCγ signaling is activated during lacrimal gland development by direct binding to Fgfr2

PLCγ1 is a large protein featuring a PIP3-binding PH domain, regulatory EF-hand and C2 domains, protein-interacting SH2 and SH3 domains, and catalytic X and Y domains (Fig. 2A) (Kadamur and Ross, 2013). Previous studies have shown that PLCγ1 recognizes a conserved phosphotyrosine residue in FGFRs (pY766 in Fgfr1 and pY769 in Fgfr2) through one of its SH2 domains, leading to phosphorylation and activation of PLCγ1 (Bae *et al*., 2009; Mohammadi *et al*., 1991). To specifically disrupt this FGFR-PLCγ1 interaction, we used CRISPR-Cas9 technology to engineer a Y769F mutation in Fgfr2 (*Fgfr2^PLCγ^*) (Fig. 2B). Offsprings were screened by restriction digestion and verified by Sanger sequencing to carry the desired Y769F mutation (Fig. 2C and D). Following backcrossing to remove potential off-target mutations, we derived MEF cells from homozygous *Fgfr2 ^PLCγ/PLCγ^*embryos. Given that MEF cells express both Fgfr1 and Fgfr2, we stimulated them with FGF9, which preferentially activates the Fgfr2 over Fgfr1. In response, the *Fgfr2 ^PLCγ/PLCγ^* MEF cells displayed robust ERK phosphorylation comparable to control cells but lacked PLCγ1^Y783^ phosphorylation (Fig. 2E). This confirms that pY769 in Fgfr2 is a critical binding site for PLCγ1, and disrupting FGF-induced PLCγ activity does not impact MAPK signaling.

**Figure 2.**
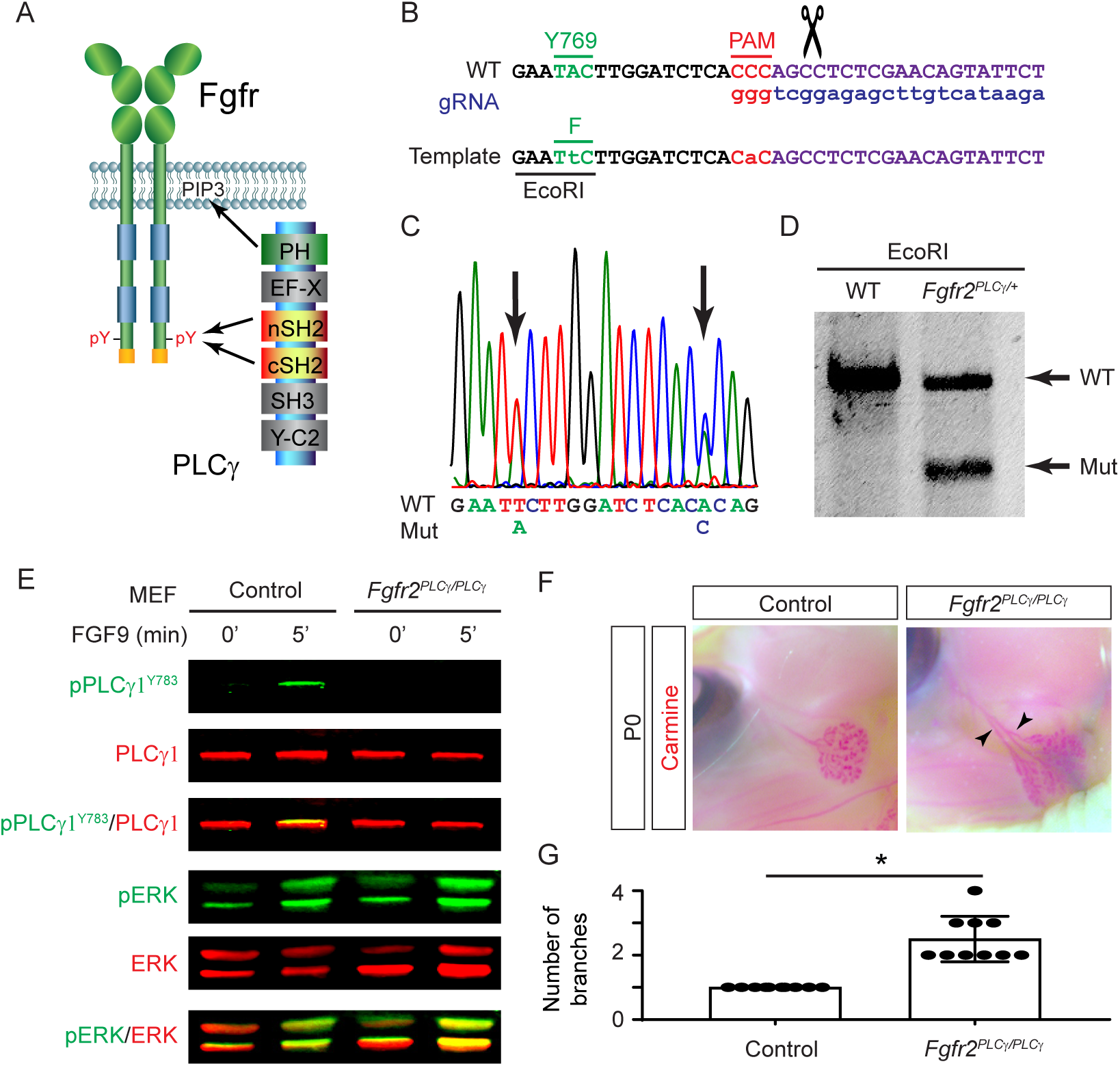
Disruption of the PLCγ binding site on Fgfr2 resulted in lacrimal gland hyperplasia. **(A)** PLCγ includes a PH domain for PIP3 interaction, EF-X and Y-C2 domains for enzymatic activity, and two SH2 and SH3 domains for protein-protein interactions, binding to the phosphotyrosine residue at the c-terminal of the FGF receptor. **(B)** Schematic diagram of CRISPR-Cas9 mediated gene targeting to generate *Fgfr2^PLCγ^* mutant. The Y769 residue of Fgfr2, essential for PLCγ binding, is altered to phenylalanine, which also introduces an EcoR1 site for genotyping. A silent mutation was introduced to disable the PAM site. **(C)** Sanger sequencing confirmed the Y769 mutation and PAM site disruption. **(D)** Genotyping PCR verified the precise targeting of the Y769F mutation. **(E)** Western blot analysis showed that FGF9 induced PLCγ1 Y783 phosphorylation in control, but not in *Fgfr2^PLCγ/PLCγ^* MEF cells, where ERK activation was unaffected. **(F)** *Fgfr2^PLCγ/PLCγ^*mutants displayed ectopic branching in the lacrimal gland (arrowheads). **(G)** Quantification of lacrimal gland branches. Student’s t-test, *n*=10, *P<0.0001.

We next examined lacrimal gland phenotype in *Fgfr2 ^PLCγ/PLCγ^*embryos. Although homozygous *Fgfr2 ^PLCγ/PLCγ^* mice are viable and fertile, carmine staining of mutant pups revealed extra branching and overgrowth of the lacrimal glands, resembling *PLCγ1* mutants (Fig. 2F and G, arrowheads). The similarity of *Fgfr2 ^PLCγ^*and *PLCγ1* mutant phenotype demonstrated that FGF signaling is the main upstream inducer of PLCγ1 in lacrimal gland development.

### Both nSH2 and cSH2 domains are critical for PLCγ1 phosphorylation and function

Given the ongoing debate concerning the roles of the two tandem SH2 domains in PLCγ1, we utilized CRISPR-Cas9 technology to generate two mouse models with targeted mutations in the essential FLVR motifs of each domain: R586A in the nSH2 (*PLCγ1^nSH2^*) and R694A in cSH2 (*PLCγ1^cSH2^*) (Bae *et al*., 2009; Huang *et al*., 2016) (Fig. 3A and B). Each line was validated through direct sequencing and restriction digestion, and subsequently crossed with the *PLCγ1^flox^* allele to extract MEF cells. In *PLCγ1^nSH2/flox^* and *PLCγ1^cSH2/flox^*MEF cells infected with control GFP-expressing adenovirus, PLCγ1^Y783^ phosphorylation was strongly stimulated 5 minutes after FGF2 treatment. After infection with Cre-expressing (Ad-Cre) to remove the *PLCγ1^flox^* allele, however, the FGF-induced PLCγ1^Y783^ phosphorylation was abolished (Fig. 3C and D). This result suggests that both SH2 domains are crucial for PLCγ1 regulation by FGFR. Furthermore, homozygous *PLCγ1^nSH2/nSH2^* and *PLCγ1^cSH2/cSH2^*mutants exhibited similar embryonic lethality and severe growth retardation at E10.5 (Fig. 3E), mirroring the phenotype of previously reported *PLCγ1* null mutants (Ji et al., 1997). These findings underscore the essential roles of both the nSH2 and cSH2 domains in PLCγ1 function in FGF signaling and embryonic development.

**Figure 3.**
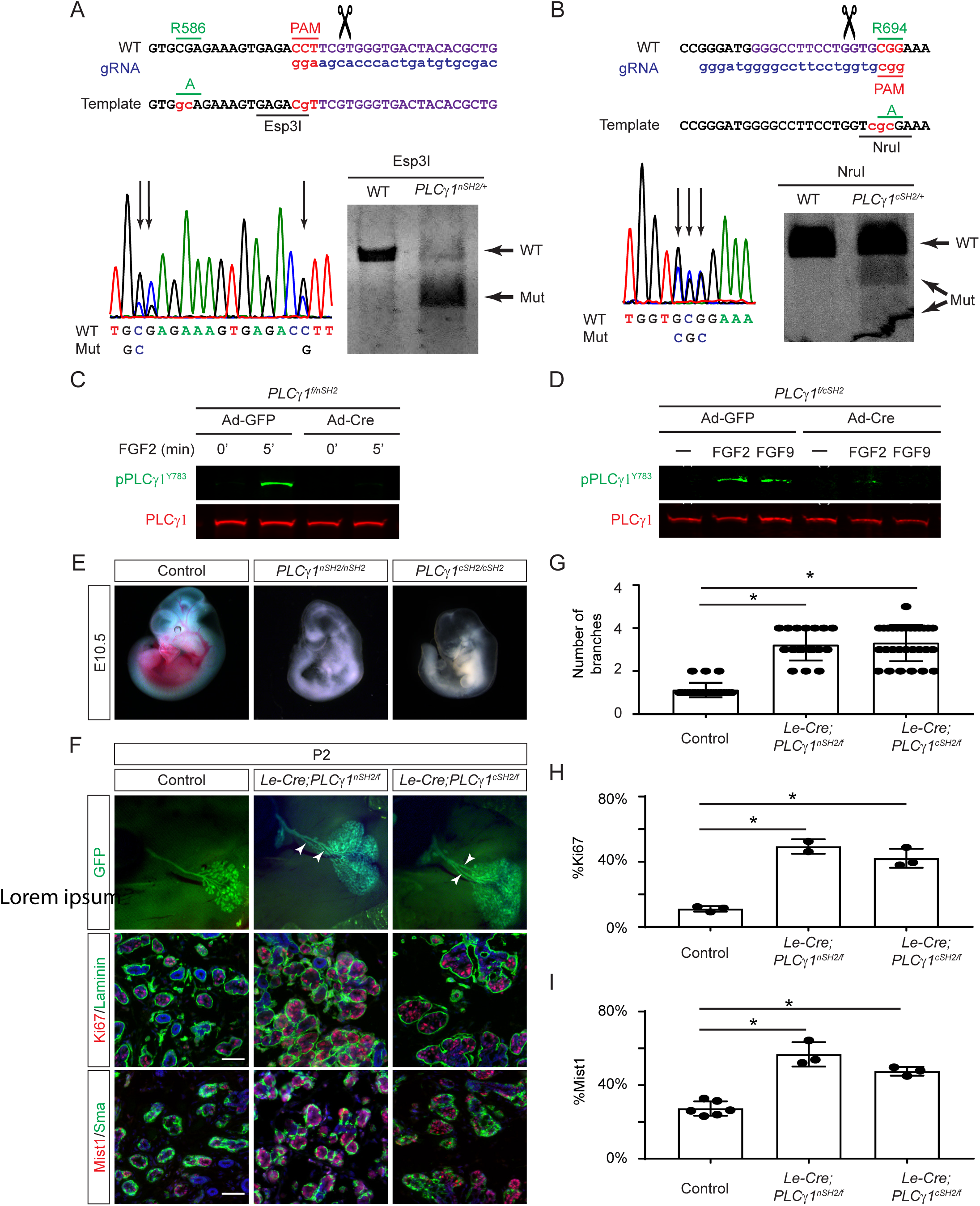
Both nSH2 and cSH2 domains are essential for PLCγ1 phosphorylation and function. **(A and B)** Using CRISPR-Cas9 gene targeting, the nSH2 domain of PLCγ1 was modified by introducing the R586A mutation and silent mutations to disrupt the PAM site and create an Esp3I restriction site in *PLCγ1^nSH2^* mutant. The cSH2 domain was altered by incorporating the R694A mutation and silent changes to introduce a NruI restriction site in the *PLCγ1^cSH2^* mutant. **(C and D)** FGF failed to induce PLCγ1 Y783 phosphorylation in either *PLCγ1^nSH2^*or *PLCγ1^cSH2^* mutant MEF cells. **(F)** Both *PLCγ1^nSH2^*and *PLCγ1^cSH2^* mutants died at E10.5 with severe growth retardation and hematopoiesis defects. **(G)** Quantification of lacrimal gland branches. One-way ANOVA, *n*>18, *P<0.0001. **(E)** Quantification of Ki67-expressing cells. One-way ANOVA, *n*>2, *P<0.001. **(F)** Quantification of Mist1-expressing cells. One-way ANOVA, *n*>3, *P<0.001. Scale bars: 50 µm.

We next generated *Le-Cre; PLCγ1^flox/nSH2^* and *Le-Cre; PLCγ1^flox/cSH2^* mice to examine lacrimal gland budding and branching morphogenesis. Consistent with the critical role of both SH2 domains for PLCγ1 function, both mutants exhibited ectopic branching similar to *Le-Cre*;*PLCγ1^flox/flox^*mutant and increasing number of Mist1 and Ki67 positive cells (Fig. 3F-I, arrowheads). These data further demonstrate that both nSH2 and cSH2 domains are critical for PLCγ1 function.

### PLCγ1 deletion does not perturb FGF-MAPK signaling in the lacrimal gland

Growth of the lacrimal gland is critically dependent on the magnitude of FGF signaling. We and others have shown that reductions in the components of FGF-MAPK signaling, including the ligand Fgf10, cofactors heparan sulfate proteoglycans and downstream effectors Pea3 transcription factors, lead to stunted lacrimal gland growth (Garg et al., 2017; Garg *et al*., 2018; Makarenkova et al., 2000; Qu *et al*., 2011; Qu *et al*., 2012). Conversely, exogenous Fgf10 can induce ectopic lacrimal gland buds, both in vivo and in explant culture (Govindarajan et al., 2000; Makarenkova *et al*., 2000). However, we did not observe any change in the pattern or intensity of *Fgf10* expression in the periocular mesenchyme (Fig. 4A, arrows). Another intriguing possibility is that PLCγ1 may regulate the trafficking of FGF receptors, as a previous study showed that FGFR1 lacking the PLCγ1-binding residue Y766 displayed reduced internalization in cell culture (Sorokin *et al*., 1994). Accordingly, deletion of *PLCγ1* may result in the accumulation of FGF receptors on the plasma membrane, which enhances FGF signaling to promote excessive lacrimal gland growth. Nevertheless, no evident changes in Fgfr2 immunostaining were observed in either E14.5 lacrimal gland buds or P0 lacrimal glands (Fig. 4A, dotted lines). Additionally, the reduction in the dosage of FGF receptors by removing a single copy of *Fgfr2* and/or two copies of *Fgfr1* in *Le-Cre*;*PLCγ1^flox/flox^*;*Fgfr2^flox/+^*and *Le-Cre*;*PLCγ1^flox/flox^*;*Fgfr1^flox/flox^*;*Fgfr2^flox/+^* mutant failed to rescue the ectopic branching phenotype (Fig. 4B and C, arrowheads). These results do not lend support to the hypothesis that *PLCγ1* deletion elevates FGF receptor levels.

**Figure 4.**
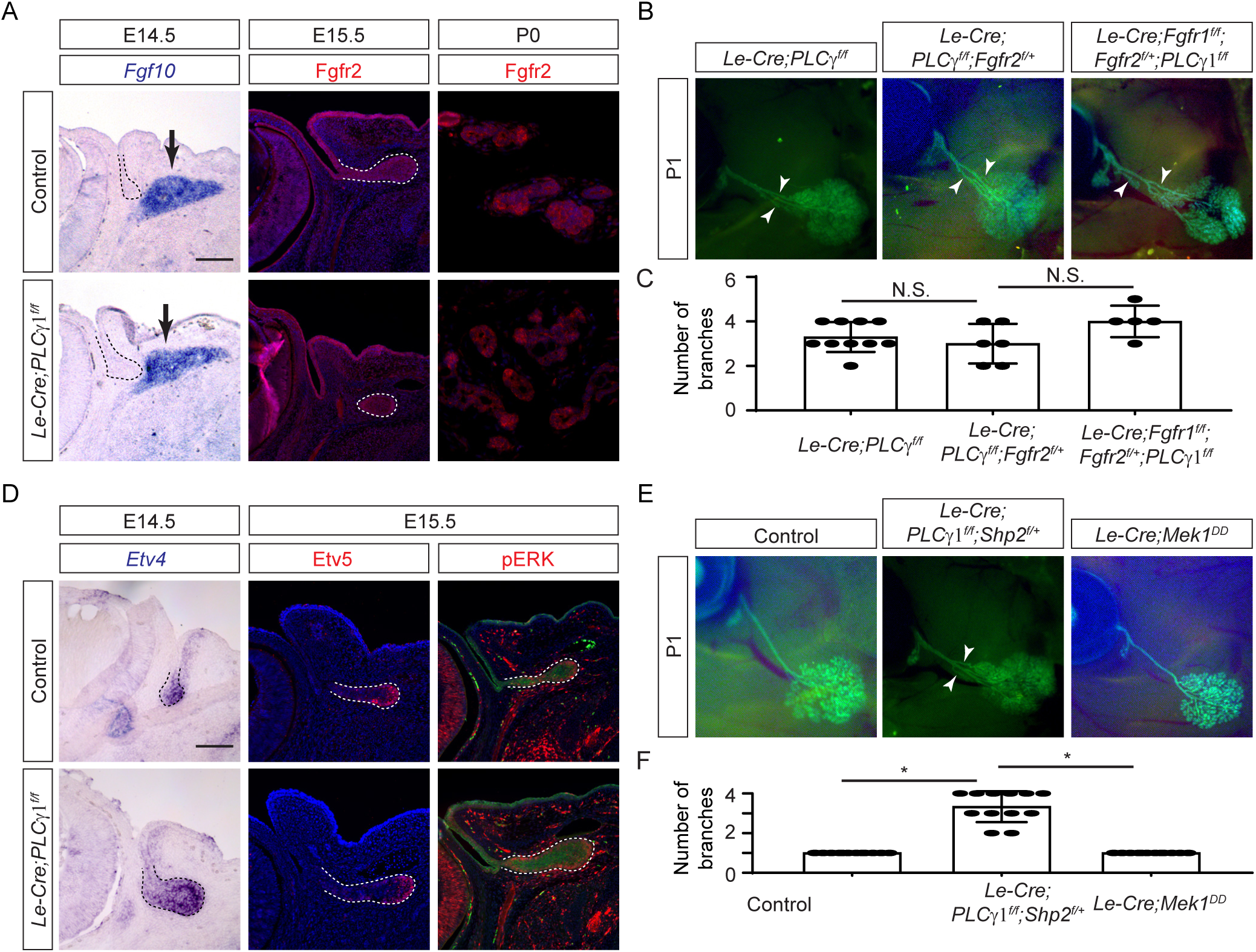
The FGF-MAPK pathway is unaffected by PLCγ1 inactivation. **(A)** Expression of *Fgf10* in the periocular mesenchyme and Fgfr2 in the lacrimal gland epithelium were unchanged in *PLCγ1* mutants. **(B)** Heterozygous deletion of *Fgfr2* and homozygous deletion of *Fgfr1* failed to rescue the ectopic branching defects in *PLCγ1* mutants (arrowheads). **(C)** Quantification of lacrimal gland branches. One-way ANOVA, *n*>5, N.S. not significant. **(D)** *PLCγ1* deletion did not affect the expression of FGF response genes *Etv4* and Etv5 and ERK phosphorylation (dotted lines). **(E)** Excessive lacrimal gland branching was not alleviated after the removal of one copy of *Shp2* in *PLCγ1* mutants or induced by activation of MAPK signaling in *Le-Cre;MEK1^DD^* animals. **(F)** Quantification of lacrimal gland branches. One-way ANOVA, *n*=12, *P<0.0001. Scale bars: 50 µm.

We further explored the possibility of altered MAPK signaling pathways contributing to the *PLCγ1* mutant phenotype. Immunostaining for pERK, a direct readout of MAPK signaling activity, and analysis of Pea3/Erm transcription factor expression, known ERK targets in lacrimal development, showed no changes in *PLCγ1* mutant glands (Fig. 4D, dotted lines) (Garg *et al*., 2017; Garg *et al*., 2018). In addition, the removal of a single copy of *Shp2*, a positive regulator of FGF signaling, failed to ameliorate *PLCγ1* mutant gland phenotype, nor did the expression of a constitutively active *MEK1^DD^* allele induce ectopic lacrimal gland branching (Fig. 4E and F, arrowheads) (Wang *et al*., 2021). Collectively, these data suggest that the FGF-MAPK pathway is not responsible for the observed phenotype in *PLCγ1* mutant lacrimal gland.

### Inactivation of PLCγ1 signaling led to sustained Akt phosphorylation upon FGF stimulation

To understand the molecular basis of the *PLCγ1* mutant phenotype, we next turned to biochemical experiments. In control *PLCγ1* ^flox/flox^ MEF cells infected with a GFP-expressing adenovirus (Ad-GFP), treatment with FGF2 led to prolonged ERK phosphorylation, but the level of pAKT peaked at 5 minutes and then returned to basal levels. In contrast, in *PLCγ1*-deficient MEF cells generated by Cre-mediated deletion (Ad-Cre), both pEKR and pAKT levels remained elevated at 30 and 60 minutes after FGF2 treatment (Fig. 5A and B). Notably, this sustained Akt phosphorylation was also observed in MEF cells carrying *PLCγ1^nSH2^* or *PLCγ1^cSH2^* mutations, further supporting PLCγ1’s role in regulating AKT signaling. Recognizing that FGF2 activates both Fgfr1 and Fgfr2, we deleted Fgfr1 in *Fgfr1^flox/flox^;Fgfr2 ^PLCγ/PLCγ^* MEF cells using Ad-Cre, which resulted in decreased ERK phosphorylation but maintained a persistent pAKT response to FGF2 stimulation. These results suggest that PLCγ1 acts downstream of the FGF receptor to temporally restrict AKT signaling to create a transient wave of activity.

**Figure 5.**
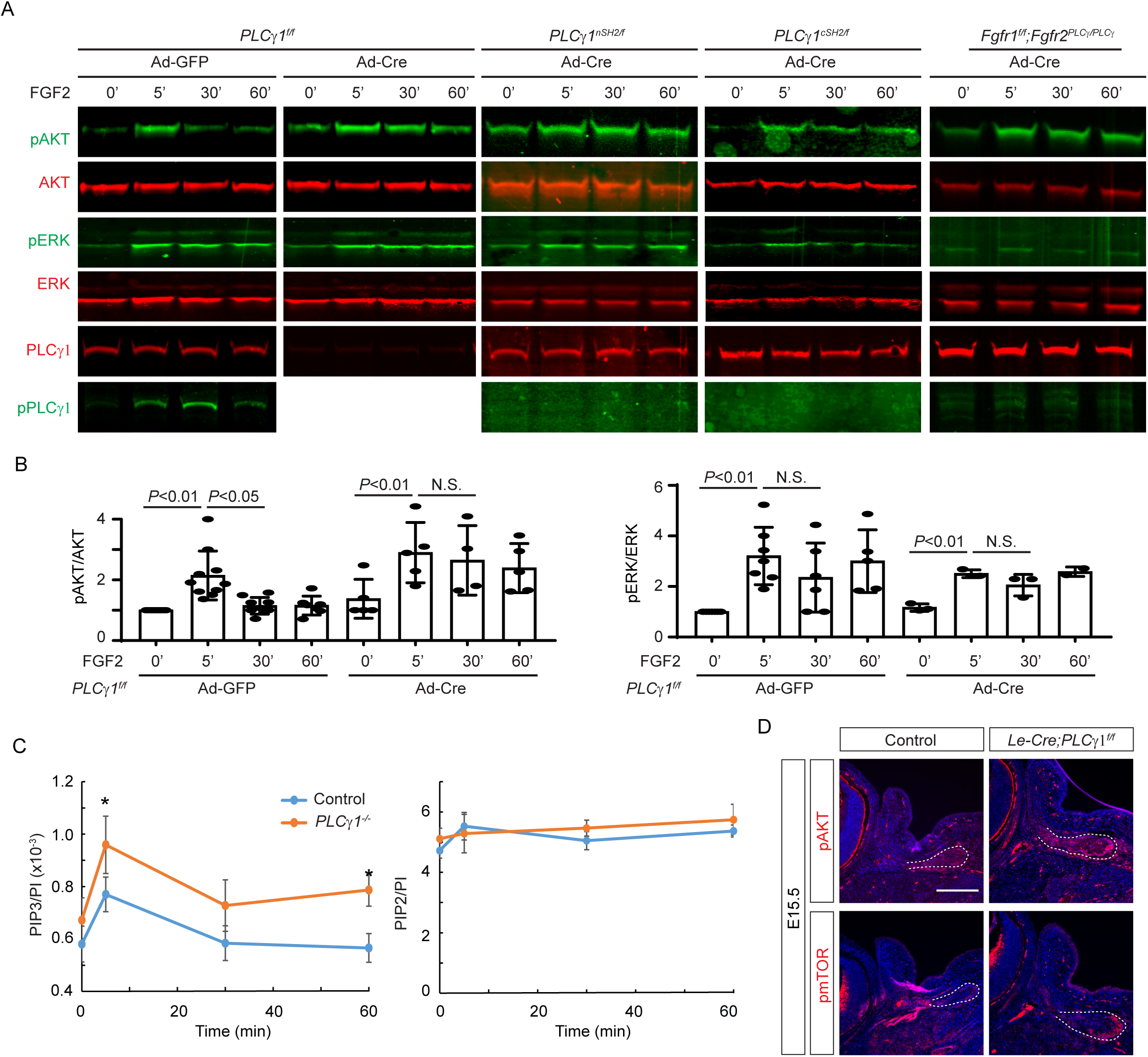
PLCγ1 inactivation resulted in sustained PI3K signaling. **(A)** AKT phosphorylation switched from a transient response peaked at 5 minutes post-FGF2 addition lasting up to 60 minutes following *PLCγ1* deletion, its nSH2 and cSH2 domains, or mutation of its binding site on Fgfr2, while ERK responses were unchanged. **(B)** Quantification of pAKT and pERK levels. One-way ANOVA, *n*>4, N.S. not significant. **(C)** Mass spectrometry measurement of the C38:4 species phosphoinositide showed that FGF2 induced higher levels of PIP3 in *PLCγ1*-deficient MEF cells compared to controls. Student’s test, *n*=3, *P<0.05. **(D)** *PLCγ1* mutant lacrimal gland buds showed stronger pAKT and pmTOR staining than controls at E15.5 (dotted lines).

To investigate how PLCγ1 influences AKT activation, we measure the levels of phosphatidylinositol 3,4,5-triphosphate (PIP3), a key activator of AKT signaling. Lipid mass spectrometry analysis showed that in MEF cells, only specific fatty acid-containing PIs (C38:4 and C38:3) showed significant increases in PIP3 levels upon FGF2 stimulation. Since the PI levels were consistent between control and *PLCγ1*-deficient MEF cells, we normalized PIP3 levels to PI levels. In line with the role of FGF in activating PI3K, there was a transient increase in the PIP3/PI ratio for the C38:4 species in control cells, peaking at 5 minutes post-FGF treatment. Crucially, *PLCγ1*-deficient cells displayed a significantly higher and more prolonged elevation in PIP3 levels, persisting up to 60 minutes after FGF2 stimulation (Fig. 5C). A similar trend was noted with the C38:3 species in all conditions, though these changes did not reach statistical significance (supplementary figure 1). The PIP2/PI ratio, however, remained unchanged in response to FGF stimulation in both control and mutant cells (Fig. 5C). These findings suggest that PLCγ1 acts as a negative regulator of FGF-AKT signaling by suppressing PIP3 production.

**Supplementary Figure 1.**
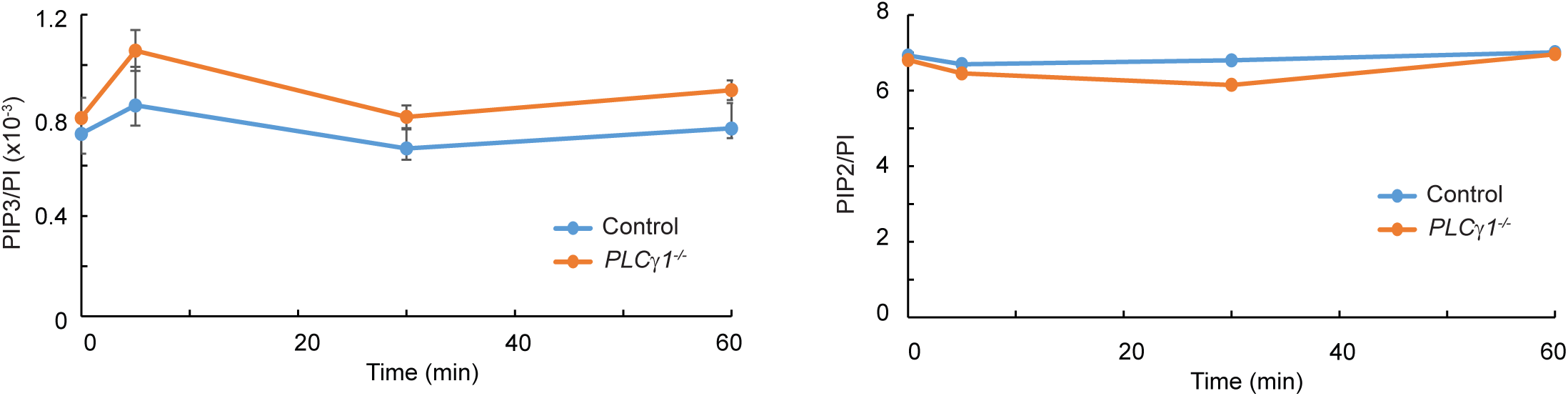
Measurement of the C38:3 fatty acid-containing phosphoinositide. The PIP3/PI ratio for the C38:3 species was elevated in *PLCγ1*-deficient MEF cells after FGF2 stimulation, but it did not reach statistical significance.

Lastly, we examined whether deletion of *PLCγ1* affected AKT signaling in vivo. At E15.5, phosphorylation of AKT and its downstream target mTOR were detected in control lacrimal gland buds (Fig. 5D, dotted lines). In contrast, the *PLCγ1* mutant buds were not only larger, but also exhibited stronger pAKT and pmTOR staining, confirming that *PLCγ1* deficiency led to increased AKT signaling in lacrimal gland development.

### Activation of the PI3K pathway leads to ectopic lacrimal gland branching

To explore the significance of sustained PIP3 synthesis in lacrimal gland development, we generated a gain-of-function model by deleting *Pten*, a lipid phosphatase that converts PIP3 back into PIP2. Notably, *Pten*-deficient (*Le-cre;Pten^flox/flox^*) lacrimal glands developed ectopic branching along the primary ducts and increased expression of the acinar marker Mist1, mirroring the phenotype seen in *PLCγ1* mutants (Fig. 6A). This led us to develop a *PLCγ1* and *Pten* double knockout model to assess their genetic interaction. Remarkably, the *Le-cre;Plcγ1^flox/flox^;Pten^flox/flox^*mutant exhibited significantly more extensive growth compared to each single mutant, with ectopic branches sprouting directly from the conjunctiva and completely obscuring the primary duct structure (Fig. 6B). In addition, the number of Mist1 cells were further amplified (Fig. 6C). These findings strongly suggest that sustained PI3K signaling, but not hyperactive MAPK signaling, can lead to ectopic lacrimal gland branching.

**Figure 6.**
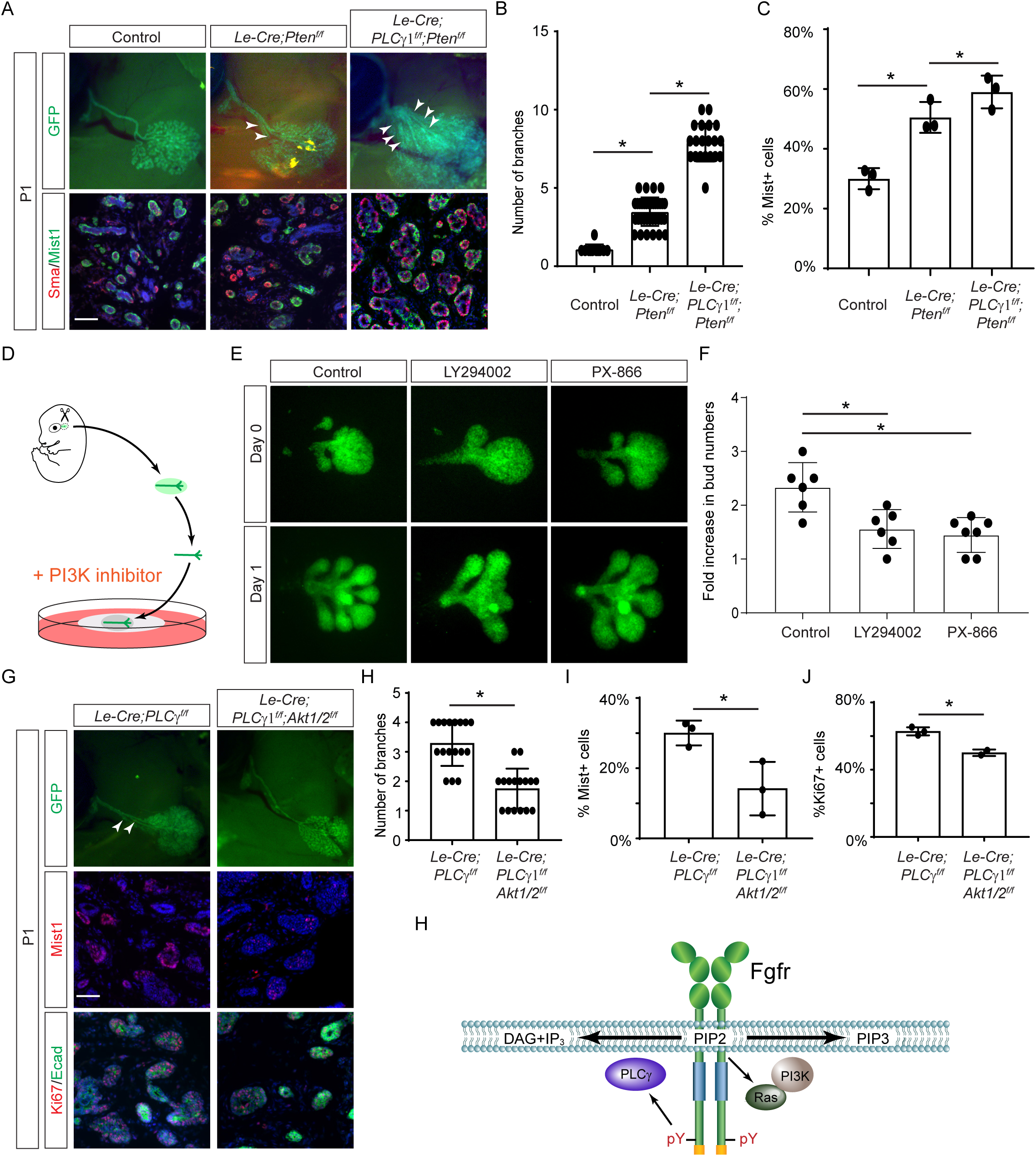
Aberrant PI3K-Akt signaling leads to branching morphogenesis defects in *the PLCγ1* mutant lacrimal gland. **(A)** Deletion of *Pten* in the lacrimal gland induced ectopic branching and further increased branching when combined with *PLCγ1* mutation (arrowheads). Additionally, there was an increase in Mist1-expressing acinar cells in both *Pten* single and *Pten*/*PLCγ1* double mutant glands. Scale bar: 50 µm. **(B)** Quantification of lacrimal gland branches. One-way ANOVA, *n*>11, *P<0.0001. **(C)** Quantification of Mist1 positive cells. One-way ANOVA, *n*=3, *P<0.01. **(D)** Schematic diagram of lacrimal gland explants. The lacrimal gland primordia were dissected from E15.5 embryos. After removal of the mesenchyme, the lacrimal gland epithelia were placed on a filter paper floating on culture media containing PI3K inhibitors (dotted lines). **(E)** After one day in culture, the lacrimal gland epithelia sprouted multiple buds, which were suppressed by PI3K inhibitors LY294002 and PX-866. **(F)** Quantification of fold changes in bud numbers. One-way ANOVA, *n*=6, *P<0.01. **(G)** Deletion of *Akt1* and *2* reduced ectopic branching in *PLCγ1* mutants (arrowheads), and decreased numbers of Mist1 and Ki67 positive cells. Scale bar: 50 µm. **(H)** Quantification of lacrimal gland branches. Student’s t-test, *n*=16, *P<0.0001. **(I)** Quantification of Mist1 positive cell. Student’s t-test, *n*=3, *P<0.05. **(J)** Quantification of Ki67 positive cells. **(B)** Crosstalk between PLCγ and PI3K signaling. FGF receptors recruits PLCγ via its phosphotyrosine residue at the C-terminal end and PI3K via Ras. The conversion of PIP2 to IP3 and DAG by PLCγ depletes the local pool of PIP2 in the vicinity of the FGF receptor, consequently limiting the ability of PI3K to generate PIP3 from PIP2. Thus, the activation of PLCγ exerts a negative regulatory effect on the PI3K pathway by reducing the availability of its substrate.

### Inhibition of PI3K/AKT signaling reversed *PLCγ1* mutant lacrimal gland phenotype

We next asked if the attenuation of PI3K signaling could reduce lacrimal gland branching. To this end, we first employed a mesenchyme-free explant model to examine directly branching morphogenesis of the lacrimal gland epithelium (Makarenkova *et al*., 2009). The lacrimal gland bud was dissected from E15.5 mouse embryos and after gentle protease treatment, separated from the mesenchyme and placed directly on top of a permeable filter before being embedded in Matrigel and floated in media (Fig. 6D). After one day of culture, the lacrimal gland primordia grew to sprout multiple branches. In explants treated with two different PI3K inhibitors, LY294002 and PX-866, the lacrimal gland expanded in size, but the number of branches was significantly reduced (Fig. 6E and F). This result showed that reducing PI3K signaling indeed inhibited lacrimal gland branching.

Lastly, we test if genetic ablation of *Akt* could rescue *PLCγ1* mutant phenotype by crossing *PLCγ1* mutants with *Akt* flox alleles. Strikingly, deletion of *Akt1* and *2* significantly reduced the number of ectopic branches in *PLCγ1* mutants (Fig. 6G and H). This was accompanied by reduced cell proliferation indicated by Ki67 staining and a fewer number of Mist1-expressing acinar cells (Fig. 6I and L). Altogether, these gain and loss of function studies demonstrated that *PLCγ1* regulated FGF-induced PI3K signaling to prevent excessive lacrimal gland branching and that the major PI3K effector responsible for these impacts was AKT.

## Discussion

PLCγ1 belongs to the Phospholipase C (PLC) enzyme family known for their roles in generating the second messengers DAG and IP3. In this study, our research revealed that PLCγ1, through its dual SH2 domains, acts as a negative regulator of FGF signaling during lacrimal gland development. Contrary to its expected effect on MAPK signaling, we showed that the deletion of *PLCγ1* resulted in the activation of AKT signaling due to a sustained increase in PIP3. This led to hyperactive branching of the lacrimal gland, which could be mimicked by deleting the PIP3 phosphatase *Pten* and mitigated by ablating *Akt*. This study demonstrates the functional significance of PLCγ and PI3K competition in vivo, highlighting a critical role for PLCγ1 in regulating branching morphogenesis.

Although PLCγ has long been recognized as a downstream pathway of FGF signaling, its precise role has remained unclear. Previous in vitro studies suggested diverse effects of disrupting the FGFR-PLCγ interaction, including reduced MAPK signaling, altered Src activity, and changes in FGF receptor internalization (Huang et al., 1995; Landgren et al., 1995; Sorokin *et al*., 1994). Notably, a Y766F mutation in Fgfr1, which abolishes PLCγ binding, caused vertebral column malformations in vivo, hinting at a negative regulatory role for PLCγ (Partanen et al., 1998). In our study, mutating the PLCγ-binding residue in Fgfr2 resulted in ectopic branching in the lacrimal gland, indicating a gain-of-function phenotype. Despite this, there were no changes in Fgfr2 levels or pERK activity in the lacrimal gland, and reducing the dosage of *Fgfr* or *Shp2* did not alter the phenotype. However, the *PLCγ1* mutant glands showed strong pAKT activity, and similar ectopic branching could be induced by deleting the PIP3 phosphatase *Pten*. Additionally, ablating AKT mitigated the aberrant phenotype of the *PLCγ1* mutant. These findings suggest that AKT, out of the large family of PIP3-effectors in mammalian cells, mediates these influences of PLCγ1 signaling on the PI3K network and lacrimal organization. The results also underscore the role of PLCγ1 as a negative regulator of FGF signaling, acting through the inhibition of PI3K-AKT signaling during lacrimal gland development.

Previous studies have established that the activation of PLCγ by receptor tyrosine kinases (RTKs) is precipitated by a cascade of molecular interactions mediated by its two SH2 domains. It was previously believed that PLCγ binds the phosphorylated tyrosine residue on RTKs via its nSH2 domain (Bae *et al*., 2009; Bunney *et al*., 2012; Hajicek *et al*., 2019). This leads to the phosphorylation of the Y783 residue on PLCγ1, which in turn competes for the cSH2 domain binding, thereby exposing the catalytic site and enabling enzyme activation. According to this framework, the nSH2 domain is essential for the PLCγ1 phosphorylation at Y783, whereas alterations in the cSH2 domain might disrupt its autoinhibitory function, as evidenced by a constitutively active human mutation in the cSH2 domain. Nonetheless, this model is contested by a recent crystallography study showing FGFR1 binding to the cSH2 domain, whereas the nSH2 domain is dispensable (Huang *et al*., 2016). Our current genetic study reveals that disrupting either nSH2 or cSH2 domains in PLCγ1 mimicked the null mutant phenotype, both in embryonic development and lacrimal gland branching. Contrary to the expected inhibitory role of the cSH2 domain, we found that the cSH2 mutation also blocked PLCγ activity and prevented FGF-induced Y783 phosphorylation. These unexpected results do not fully agree with either model of the PLCγ activation mechanism, suggesting that further studies are needed to elucidate the structure and function of this enzyme.

PLCγ and PI3K are key enzyme families in phospholipid metabolism, yet it’s unclear if their competition for the substrate PIP2 leads to crosstalk between their pathways. The relative scarcity of PIP3—typically less than 5% of PIP2—might suggest that even substantial PIP2 hydrolysis by PLCγ leaves sufficient substrate for PI3K to generate PIP3 (Balla, 2013; Delage et al., 2013). However, the rapid turnover and low abundance of PIP2 (less than 1% of total phospholipids) raise the possibility that prolonged PLCγ activation could deplete PIP2 to levels that hinder PIP3 synthesis. Supporting this idea, our cell culture studies found that deletion of *PLCγ1* did not alter AKT phosphorylation until 30 minutes post-FGF stimulation, indicating a relatively slow influence of PLCγ1 on PI3K activity. Surprisingly, lipid mass spectrometry in *PLCγ1*-deficient MEF cells showed a significant reduction in PIP3, but not PIP2. It is possible that PLCγ1 signaling causes a reduction in FGF-stimulated PIP3 accumulation by inhibition of PI3K activation or augmentation of PIP3 phosphatase activity-although the ability of the *Pten* deletion to synergize with *PLC*γ1 deficiency makes this latter concept less likely. Alternatively (but not mutually exclusively), a model of localized PIP2 depletion, within specialized membrane microdomains like lipid rafts that act as signaling hubs harboring FGF receptors (Hilgemann, 2007), could apply. In these confined spaces where both PI3K and PLCγ1 are recruited, the limited diffusion and replenishment of PIP2 could lead to localized depletion significantly impacting PIP3 production, even though the total cellular PIP2 appears unchanged. Given that PI3K also activates PLCγ through PIP3-mediated targeting of the PLCγ PH domain, this mechanism of shifting phospholipid metabolism forms an essential part of the delayed negative feedback loop that may shape the dynamics of both PLCγ and PI3K signaling across various RTK pathways.

## Materials and methods

### Mice

*Akt1^flox^* and *Akt2^flox^* mouse were kindly provided by Drs. Rebecca Haeusler (Columbia University) and Paul M. Titchenell (University of Pennsylvania) (Leavens et al., 2009; Wan et al., 2012), *Fgfr2^flox^* by Dr. David Ornitz (Washington University Medical School, St Louis, MO) (Yu et al., 2003), *Le-Cre* by Richard Lang (Children’s Hospital Research Foundation, Cincinnati, OH) (Ashery-Padan et al., 2000), *Plcγ1^flox^* by Renren Wen and Demin Wang (Versiti Blood Research Institute, Milwaukee, WI) (Fu et al., 2010), *Shp2^flox^* by Gen-sheng Feng (UCSD, San Diego, CA) (Zhang et al., 2004). *Fgfr1^flox^* (Stock No: 007671), *R26R-LSL-Mek1^DD^*(Stock No: 012352) and *Pten^flox^* (Stock No: 006440) were obtained from the Jackson Laboratory (Bar Harbor, ME, USA). Animals were maintained on mixed genetic backgrounds. In all conditional knockout experiments, mice were maintained on a mixed genetic background and *Le-Cre* only or *Le-Cre* and heterozygous flox mice were used as controls. All procedures were conducted in compliance with the protocols approved by the Institutional Animal Care and Use Committee of Columbia University.

*Fgfr2^Y788F^* (*Fgfr2^PLCγ^*), *Plcγ1^R586A^* (*PLCγ1^nSH2^*) and *Plcγ1^R694A^* (*PLCγ1^cSH2^*) mice were developed using the modified Easi-CRISPR technique (Quadros et al., 2017). This involved identifying highly specific gRNAs close to the target mutation sites using online tools (http://crispor.tefor.net), which were chemically synthesized by IDT with modifications such as 2’-O-methyl 3’phosphorothioate and end-blocking Alt-R. Single-stranded donor templates featuring the necessary amino acid changes and disrupted PAM sites, along with silent mutations to create restriction enzyme sites, were injected into mouse zygotes along with a pre-assembled gRNA-Cas9 enzyme complex at Columbia University Medical Center’s transgenic facility. Founder animals with the correct genetic modifications were confirmed through direct sequencing. The gRNAs used included TCTTATGACAAGCTCTCCGACCC for *Fgfr2^Y788F^* (*Fgfr2^PLCγ^*), CAGCGTGTAGTCACCCACGAAGG for *Plcγ1^R586A^* (*PLCγ1^nSH2^*), and GGGATGGGGCCTTCCTGGTG CGG for *Plcγ1^R694A^* (*PLCγ1^cSH2^*).

### Visualization of the whole mount lacrimal gland

For embryos possessing the *Le-Cre* transgene, their lacrimal glands were visualized using the GFP signal under a Leica MZ16F fluorescence dissecting microscope. For embryos lacking GFP, the lacrimal glands were stained using aceto-carmine as previously described (Pan *et al*., 2008). The procedure began with the decapitation of the embryos, followed by dissection to expose the lacrimal gland, which was then fixed in 4% paraformaldehyde (PFA) overnight. Post fixation, the heads were dehydrated in 70% ethanol and subsequently immersed in a 0.5% carmine solution (C-1022, Sigma, St. Louis, MO) prepared in 45% boiling acetic acid for 5-10 minutes. The lacrimal gland was then destained using a sequence of 70% ethanol for three minutes, 1% acid alcohol (1% HCl in 70% ethanol) for two minutes, and 5% acid alcohol (5% HCl in 70% ethanol) for one minute. The glands were finally examined under a Leica MZ16F dissection microscope.

### Immunohistochemistry and RNA in situ hybridization

Histology and immunohistochemistry were performed on the paraffin and cryosections as previously described (Carbe et al., 2012; Carbe and Zhang, 2011). RNA in situ hybridization and immunostaining were performed on the cryosections (10 µm) (Carbe et al., 2013). Antibodies used are Sox9 (#82630), Sox10 (#78330), FGFR2 (#23328), Mist1 (#14896), phospho-ERK1/2 (#4370), phospho-Akt (#4060), and phospho-mTOR (#2481) from Cell Signaling Technology, PLCγ1 (#610028), Ki67 (#550609) and E-cadherin (#610181) from BD Pharmingen, α-SMA (#C6198) and Laminin (L9393) from Sigma-Aldrich, Pax6 (#RPB-278P) from Biolegend. Immunostaining of phospho-ERK1/2, phospho-Akt, and phospho-mTOR was done using a Tyramide Signal Amplification kit (PerkinElmer Life Sciences, Waltham, MA). The in situ probes used were *Fgf10* (IMAGE clone, 6313081) (from Open Biosystems, Huntsville, AL, USA) and *Etv4* (from B. Hogan, Duke University Medical Center, Durham, NC, USA).

### MEF cells and Western blot

Primary mouse embryonic fibroblast (MEF) cells were derived from E13.5 embryos and cultured in Dulbecco’s modified Eagle’s medium (DMEM) supplemented with 10% fetal bovine serum. MEF cells carrying flox genes were infected with adenovirus expressing Cre recombinase (Ad5-cre) or GFP (Ad5-GFP) (Gene Transfer Vector Core, University of Iowa, IA) for 5 days to achieve gene deletion. Following a 16-hour starvation period, cells were treated with FGF2 (50ng/ml), FGF9 (50ng/ml) at various time intervals. Subsequently, the cells were washed with cold PBS and lysed in ice-cold CelLytic reagent (C2978, Sigma-Aldrich) for protein extraction. Proteins were then analyzed by Western blot using a standard protocol. The antibodies employed included ERK1/2 (#4695), Akt (#2920), Phospho-Akt (Ser473) (#4060) and phospho-PLCγ1 (#14008) from Cell Signaling Technology, along with PLCγ1 (#610028) from BD Transduction and phospho-ERK1/2 (sc-7383) from Santa Cruz Biotechnology.

### Lipid Mass Spectrometry

1.5-2×10^5^ primary mouse embryonic fibroblast (MEF) cells were seeded in 35 mm dishes and treated with adenovirus and FGF2 as described above. To terminate the stimulation, cells were washed with cold PBS and lysed in 600 µl of ice-cold 1M HCl, then collected into 2ml safe-lock Eppendorf tubes on ice. The samples were centrifuged at 14,000 rpm at 4 °C for 10 minutes. The supernatant was removed, and the pellet was immediately snap-frozen in liquid nitrogen. The major phosphoinositide species, including C34:1, C34:2, C36:1, C36:2, C38:3, and C38:4, were quantified using mass spectrometry essentially as previously described (Clark et al., 2011), using a QTRAP 4000 (AB Sciex) mass spectrometer and employing the lipid extraction and derivitisation method described for adherent cells, with the modification that 10 ng C17:0/C16:0 PI ISD were added to primary extracts. To compensate for potential losses of lipids during extraction and analysis, a mixture of PI, PIP2, and PIP3 internal standards (ISD) was added to each sample at the start of the extraction process. As per standard protocol, the response of the endogenous lipids was normalized to the response of the corresponding ISD, providing a ratio indicative of the specific lipid levels. To control for variations in cell input, the response ratio of PIP2 or PIP3 was further normalized by the PI response ratio. Three replicates were prepared for each experimental condition.

### Lacrimal gland epithelium explant culture

Lacrimal glands expressing the *Le-Cre* GFP reporter were dissected from E15.5 mouse embryos and processed as previously described (Makarenkova *et al*., 2009). The glands underwent enzymatic dissociation with a Trypsin Pancreatin solution (2.25% Trypsin (T4799, Sigma), final 0.75% Pancreatin (P3292, Sigma) in Mg^2+^-, Ca^2+^-Free HBSS) trypsin (2.25%, T4799, Sigma) and pancreatin (0.75%, P3292, Sigma) solution on ice for 35 minutes. Subsequently, the glands were transferred to DMEM supplemented with 10-15% FBS, where the mesenchyme was manually removed using BD PrecisionGlide needle (26G x ½). The isolated lacrimal gland epithelial explants were then embedded in growth factor reduced Matrigel on a floating insert (MF-Millipore HABP02500) and cultured on top of CMRL media (Invitrogen #21540026) containing 10% FBS, Antibiotics and Antimycotics, supplemented with either 250 µM LY294002 (#9901) or 3 µM PX866 (#13055), both from Cell Signaling Technology. After incubating for one day at 37°C in a 5% CO2 atmosphere, the explants were examined to evaluate the branching patterns of the GFP-positive lacrimal gland epithelium.

### Quantification and Statistical Analysis

The relative lacrimal gland sizes were measured using Image J and normalized against the control. The number of lacrimal gland branches before reaching the junction of the supraorbital and infraorbital branches of the stapedial artery were counted. The proportions of Mist1, Ki67, and TUNEL positive cells were calculated as percentages of the total number of DAPI-positive cells within each epithelial cluster. The values of PIP3 and PIP2 were normalized against PI. Statistical analyses were conducted using GraphPad Prism 7 software. Sample sizes were not predetermined prior to the experiments. Results are presented as mean ± standard deviation (s.d.). An unpaired two-tailed t-test was employed for comparisons between two groups, while a one-way ANOVA with Tukey’s multiple comparisons test was used to analyze differences among three or more groups.

## Acknowledgments

The authors thank Drs. Ruth Ashery-Padan, Gen-sheng Feng, Rebecca Haeusler, Richard Lang, David Ornitz, Paul M. Titchenell, Renren Wen and Demin Wang for mice, Bridgit Hogan for reagent, Helen Makarenkova for advice on explant culture. The work was supported by NIH grants (R01EY018868 and R01EY031210 to X.Z. and R01EY034451 to C.T.) and BBRSC Institute Strategic Programme Grant (BB/Y0006925/1 to L.S). Q.W. is supported by a Pathway to Independence Award (K99EY032171). The Columbia Ophthalmology Core Facility is supported by NIH Core grant 5P30EY019007 and unrestricted funds from Research to Prevent Blindness (RPB).

